# Down-regulated GAS6 impairs synovial macrophage efferocytosis and promotes obesity-associated osteoarthritis

**DOI:** 10.1101/2022.09.20.508661

**Authors:** Yao Zihao, Qi Weizhong, Liu Liangliang, Shao Yan, Zhang Hongbo, Yin Jianbin, Pan Haoyan, Guo Xiongtian, Liu Anling, Cai Daozhang, Bai Xiaochun, Zhang Haiyan

## Abstract

Obesity has always been considered a significant risk factor in OA progression, but the underlying mechanism of obesity-related inflammation in OA synovitis remains unclear. The present study found that synovial macrophages infiltrated and polarized in the obesity microenvironment and identified the essential role of M1 macrophages in impaired macrophage efferocytosis using pathology analysis of obesity-associated OA. The present study revealed that obese OA patients and ApoE^−/−^ mice showed a more pronounced synovitis and enhanced macrophage infiltration in synovial tissue, accompanied by dominant M1 macrophage polarization. Obese OA mice had a more severe cartilage destruction and increased levels of synovial apoptotic cells than OA mice in the control group. Enhanced M1-polarized macrophages in obese synovium decreased growth arrest-specific 6 (GAS6) secretion, resulting in impaired macrophage efferocytosis in synovial apoptotic cells. Intracellular contents released by accumulated apoptotic cells further triggered an immune response and lead to a release of inflammatory factors, such as TNF-α, IL-1β, and IL-6, which induce chondrocyte homeostasis dysfunction in obese OA patients. Intra-articular injection of GAS6 restored the phagocytic capacity of macrophages, reduced the accumulation of local apoptotic cells, and decreased the levels of TUNEL- and caspase-3-positive cells, preserving cartilage thickness and preventing the progression of obesity-associated OA. Therefore, blocking M1 macrophage polarization or intra-articular injection of GAS6 is a potential therapeutic strategy for obesity-associated OA.

## Introduction

Osteoarthritis (OA) is a common, chronic, degenerative joint disease and a significant cause of joint pain and even disability.^1^ Epidemiological investigations have documented that obesity is one of the significant risk factors for OA.^2,3^ A meta-analysis of joint replacements in obese patients in 2010 has shown that the risk of knee OA was five times higher in obese patients than in healthy individuals.^4^ At the same time, being “overweight” (i.e., obesity) doubled the proportion of joint replacement treatments required later in life.^5^ At present, the incidence of obesity continues to increase.^6^ It is estimated that by 2025, the global incidence of obesity will reach 18% in men and 21% in women.^7^ Thus, elucidating the mechanisms by which obesity promotes OA development is essential for OA prevention and treatment.

It was initially believed that obesity affects OA by changing certain mechanical factors. However, the progression of OA continues in non-weight-bearing areas, even after the line of the force is corrected.^8,9^ Moreover, these obesity-related mechanical factors cannot be justified in the development of OA in non-weight-bearing joints such as the hands.^10,11^ Scientists have paid specific attention to the effects and roles of various pro-inflammatory cytokines and adipokines in obesity.^12–14^ Clinical data have confirmed that obese OA patients are often associated with severe chronic synovitis, which plays an essential role in the pathogenesis and progression of OA.^15,16^ Our previous findings have indicated that synovial macrophage polarization is significantly correlated with synovitis in OA progression.^17–19^ When synovitis occurs, macrophages are stimulated by various cytokines and consequently release inflammatory mediators.^20^ At the same time, excess energy caused by obesity leads to changes in various cell functions, including angiogenesis and inflammatory cell infiltration.^21^ However, the effect of obesity on synovial hyperplasia and macrophage polarization in OA development has not been reported yet. We speculate that changes in tissues and organs caused by obesity may also affect synovitis, which in turn affects the process of OA.

GAS6 is a secreted glycoprotein widely expressed throughout the body.^22,23^ It is well known for its vital role in bridging phosphatidylserine on the surface of apoptotic cells with its receptors Tyro3, Axl, and Mer, triggering the engulfment of apoptotic cells in an inflammatory environment.^22^ This macrophage-related phagocytic process is also known as “efferocytosis,” which is beneficial for resolving inflammation.^23^ Prior studies have shown that impaired efferocytosis weakens the ability to clear apoptotic cells, inducing the release of inflammatory factors and ultimately causing synovitis.^24,25^ Nevertheless, obesity-related macrophage polarization and the effect of obesity on efferocytosis remain unclear.

The present study found that obese OA patients and obese ApoE^−/−^ mice are more prone to M1 macrophage infiltration in synovial tissue. Obese OA mice had more severe cartilage destruction and increased synovial apoptotic cells than OA mice in the control group. Down-regulation of GAS6 by M1 macrophages resulted in impaired efferocytosis in synovial apoptotic cells, causing synovial hyperplasia and obesity-associated OA development. These data suggested that targeting macrophage phagocytosis and polarization in obese patients with OA may be a potential therapeutic strategy.

## Results

### 1. Synovial tissues are highly hyperplastic in obese OA patients and infiltrated with more polarized M1 macrophages than non-obese OA patients

To investigate the role of obesity in synovial tissue in OA patients, levels of total cholesterol, triglycerides, and body mass index (BMI) were examined in different patients. All subjects were divided into the following four groups based on the obtained values for these three factors: normal healthy individuals, normal OA patients, obese patients, and obese OA patients (Supplementary Table 1). Consistent with our previous study, highly hyperplastic synovial tissues and abundant inflammatory cell infiltration were observed in human OA synovial tissue, combined with a significantly higher synovitis score than normal controls. Interestingly, the synovium tended to be hyperplastic in obese patients and reached a maximum in obese OA patients among the four groups (Fig. 1A and C).

**Figure 1.**
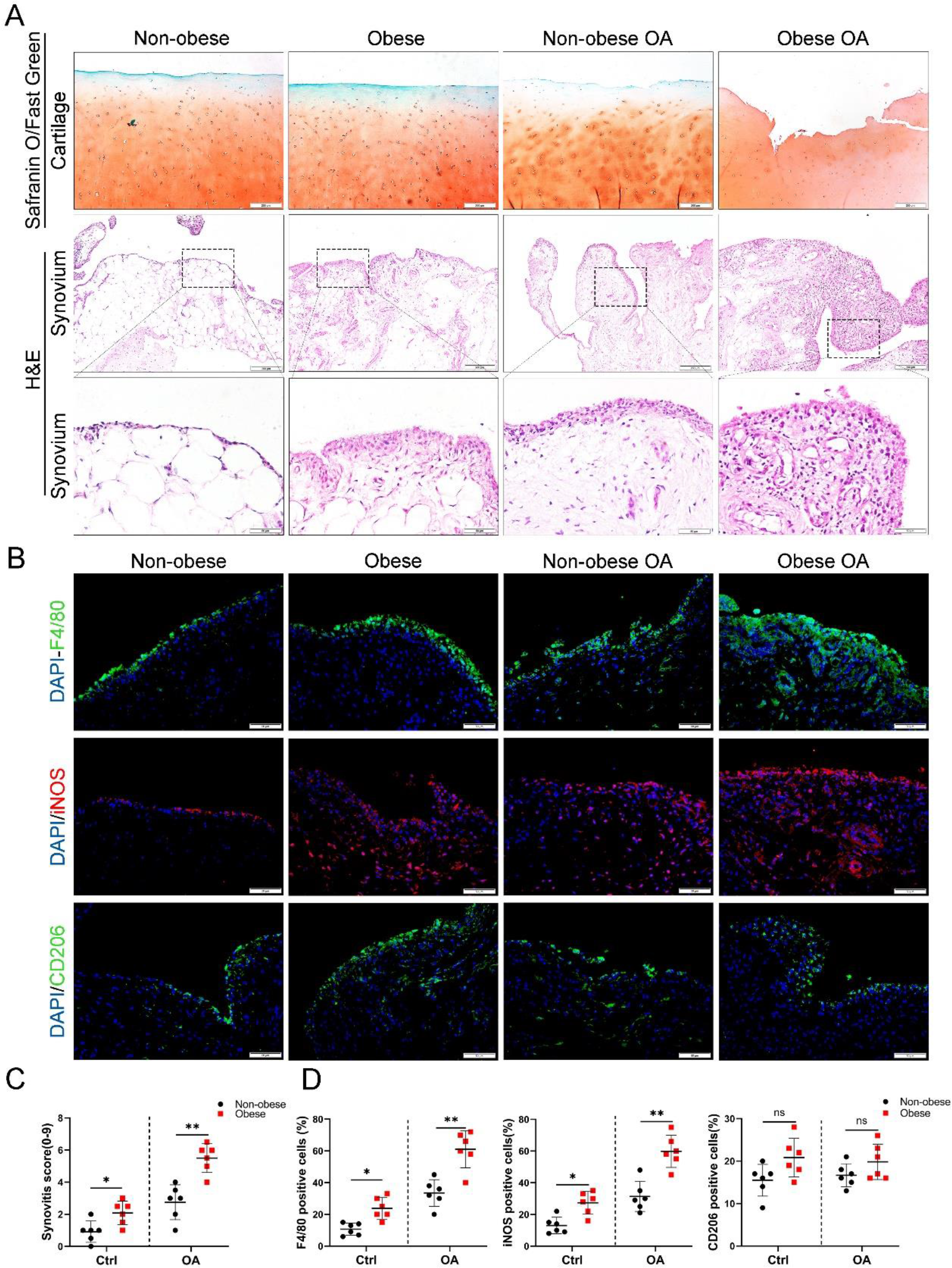
Synovial hyperplasia and macrophage polarization in obese OA patients. (A) Safranin O and Fast Green staining (top) of human articul.ar cartilage, H&E staining (lower) of synovial tissue from normal, normal osteoarthritic, normal obesity, and obesity-associated OA samples. Scale bar: 200 μm, 50 μm; (B) Immunofluorescence of F4/80, iNOS, and CD206 in normal and OA synovial tissues from non-obese and obese patients. F4/80: green; iNOS: red; DNA: blue; Scale bar: 500 μm. (C) Quantification of synovitis score in normal, normal OA, obese, and obese OA human synovial tissue samples; (n=6 per group); (D) Quantification of F4/80, iNOS, and CD206-positive macrophages as a proportion of total lining cell population in (B). *P<0.05; **P<0.01, ***P<0.001, NS=not significant. One-way analysis of variance (ANOVA) was performed. Data are shown as mean ± SD.

We further investigated the polarization level of macrophages in synovial tissues by staining with F4/80 (macrophage markers), inducible nitric oxide synthase (iNOS; M1 macrophage marker), and CD206 (M2 macrophage marker). As a result, the number of M1 macrophages in the synovial tissue of the OA group increased significantly compared to the control group. Moreover, synovial tissue in obese OA patients was infiltrated with more M1 macrophages than that in non-obese OA patients (Fig. 1B and D). These results indicate that synovial tissues were highly hyperplastic in obese OA patients and infiltrated with more polarized M1 macrophages than in non-obese OA patients.

### 2. Obesity promotes synovial M1 macrophage accumulation, synovitis, and OA development in mice

The ApoE^−/−^ mouse model was established to further explore the role of obesity in OA development, as it is considered an ideal model for investigating obesity. The body weight and plasma lipid levels were markedly elevated in ApoE^−/−^ mice after administering a high-fat and high-energy diet (Supplementary Tables 2 and Table 3). There were no significant differences in knee OA OARSI scores between ApoE^−/−^ and C57BL/6 mice four weeks post-surgery (Supplementary Fig. S1A and B). However, the OARSI score was significantly elevated in ApoE^−/−^ mice eight weeks post-surgery (Fig. 2A and B), accompanied by a higher synovitis score and more infiltrated inflammatory cells (Fig. 2A and C), indicating that obesity may promote OA development in mice. In addition, ApoE^−/−^ OA mice expressed less aggrecan on cartilage and more MMP13 on cartilage and synovium than C57BL/6 mice (Fig. 3D and E). Notably, the percentage of M1-like macrophages was increased with OA progression and reached a maximum in obese ApoE^−/−^ OA mice eight weeks post-surgery. However, the proportion of positive cells for M2-like macrophages in the OA synovium showed no significant change at both 4 and 8 weeks post-surgery (Fig. 2F and G, Supplementary Fig. S1C and D). These findings suggest that obesity exacerbates synovitis and M1-polarized macrophage accumulation during OA progression in mice.

**Figure 2.**
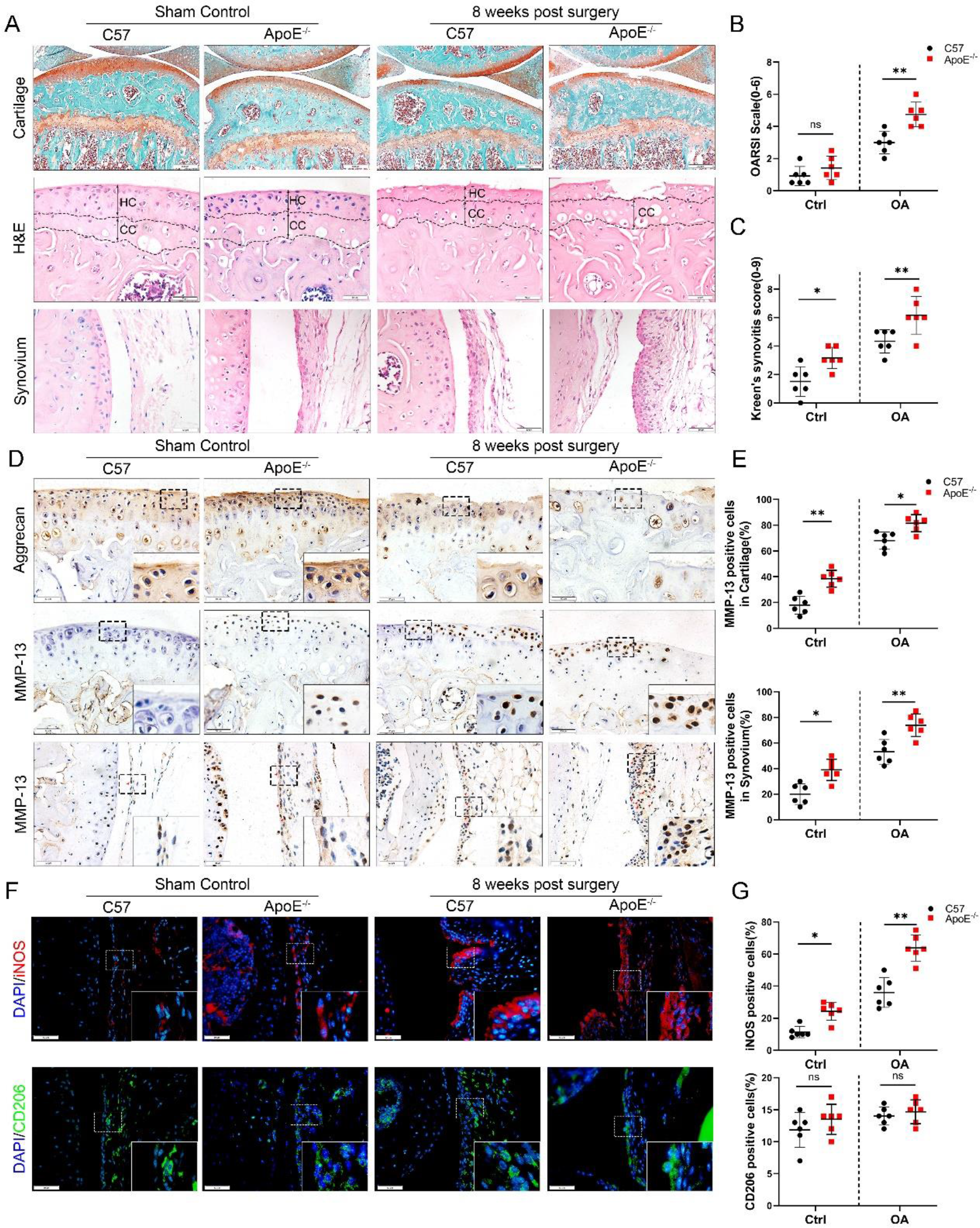
Cartilage loss, synovial hyperplasia, and macrophage polarization in ApoE^−/−^ OA. (A) Safranin O and Fast Green (first line) and H&E (second line) staining of controls and DMM knee cartilage or synovial membrane from normal and ApoE^−/−^ mice. Scale bar: 200 μm, 50 μm; (B) Quantitative analysis of Osteoarthritis Research Society International (OARSI) scale in A (second line), n=6 per group. (C) Synovitis score for joints described in (A) (third line), n=6 per group. (D) Immunohistochemical staining for aggrecan (first line) and MMP-13 (middle and bottom) in controls and DMM knee cartilage from normal and ApoE^−/−^ mice. Scale bar: 50 μm; (E) Quantification of MMP13-positive cells from cartilage or synovium in (D), n=6 per group. (F) Immunofluorescence staining for iNOS (first line) and CD206 (second line) in controls and DMM synovial tissues from normal and ApoE^−/−^ mice. Scale bar: 50 μm; (G) Quantification of iNOS- and CD206-positive cells as a proportion of lining cell population in (F), n=6 per group. *P<0.05; **P<0.01, NS=not significant. One-way analysis of variance (ANOVA) was performed. Data are shown as mean ± SD.

**Figure 3.**
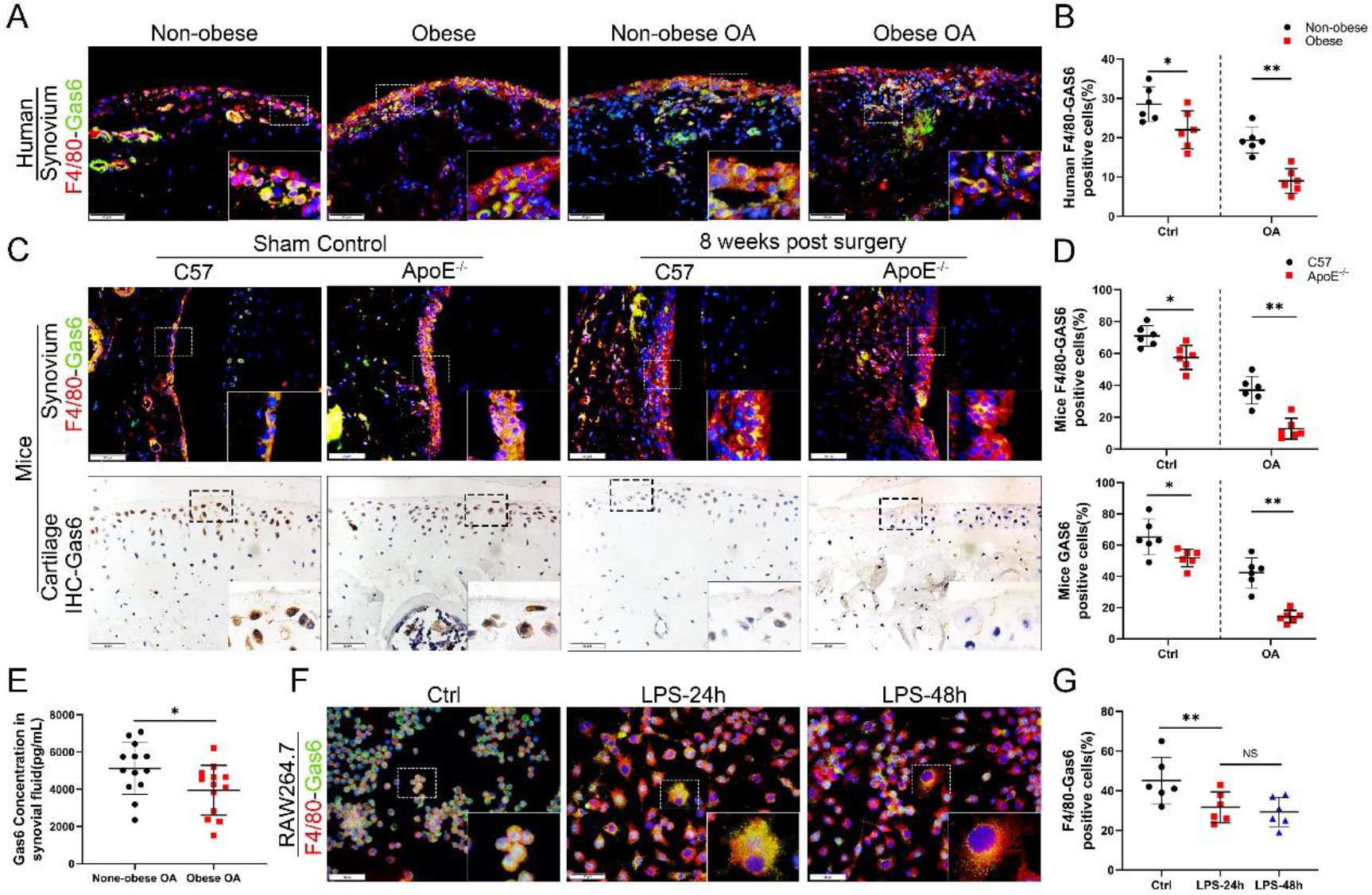
Loss of GAS6 expression in chondrocytes and synoviocytes in obese OA patients and ApoE^−/−^ OA mice. (A) Immunofluorescence staining for F4/80 (red) and GAS6 (green) in synovial tissue from non-obese, non-obese OA, obese, and obese OA patients. Scale bar: 50 μm. (B) Quantification of F4/80-GAS6-positive macrophages as a proportion of total lining cell population in (A), n=6 per group. (C) Immunofluorescence staining (first line) for F4/80 (red) and GAS6 (green) in synovial tissue and immunohistochemical staining of GAS6 (second line) in cartilage of controls and DMM from C57BL/6 and ApoE^−/−^ mice. Scale bar: 50 μm. (D) Quantification of F4/80-GAS6 positive macrophages (yellow) as a proportion of total F4/80 positive cells in (C) (first line). Quantification of GAS6-positive cells in (C) (second line), n=6 per group. (E) Enzyme-linked immunosorbent assay (ELISA) for GAS6 in synovial fluid of non-obese and obese OA patients, n=13 per group. (F) Immunofluorescence staining for F4/80(red) and GAS6 (green) in RAW264.7 cells treated with LPS for 24 and 48 h. Scale bar: 50 μm; (G) Quantification of F4/80-GAS6-positive macrophages (yellow) as a proportion of total F4/80-positive cells (red), n=6 per group. * P<0.05, ** P<0.01, *** P<0.001, NS=not significant. One-way analysis of variance (ANOVA) was performed. Data are shown as mean ± SD.

### 3. GAS6 expression is inhibited in synovial macrophages during obesity-associated OA development

The link between M1 macrophages and synovial hyperplasia during obesity-associated OA progression was further explored. GAS6, a member of the vitamin K-dependent protein family, has been previously found to be down-regulated in LPS-induced bone marrow-derived macrophages (BMDMs) compared to the controls (GSE53986, Supplementary Fig. S2A, Supplementary Table 4).^26^ GAS6 was investigated as a critical factor regulating cell proliferation and apoptosis by binding to its receptor Axl. The GAS6/Axl effects on OA remain unclear. The present study explored the association between synovial macrophage polarization types and the GAS6/Axl pathway in obesity-associated OA. Immunofluorescence staining analysis indicated that macrophage release of GAS6 expressed in both human and mouse normal synovial tissues tended to be diminished, especially in obesity-associated OA (Fig. 3A–D). As cartilage loss increased, the expression of GAS6 in chondrocytes also decreased significantly (Fig. 3C and D). Moreover, ELISA revealed a significant decrease in GAS6 levels in the synovial fluid from obese OA patients than in non-obese individuals (Fig. 3E). An *in vitro* study in polarized M1 macrophages enhanced by LPS confirmed the M1 macrophage-associated reduction of GAS6 and increases in IL-1, IL-6, and TNF-ɑ (Fig. 3F and G, Supplementary Fig. S2B and C). Interestingly, F4/80-Axl double staining revealed no significant difference in the number of Axl-positive cells in the synovium of obese OA patients, obese ApoE^−/−^ OA mice, and control subjects (Supplementary Fig. S2D–G). These results indicate a potential role of M1 macrophage-mediated GAS6 in obesity-associated OA development.

### 4. GAS6 is involved in obesity-mediated inhibition of macrophage efferocytosis

Efferocytosis is an indispensable process through which dead and dying cells are removed by phagocytic cells. Immunochemical TUNEL and caspase-3 staining in the present study revealed that the number of apoptotic cells was increased in synovial tissues with OA progression, which increased significantly in obese ApoE^−/−^ OA and obese OA patients (Fig. 4A–D). Previous studies have shown that efferocytosis of apoptotic cells induced by macrophages is impaired in inflammatory diseases. Still, its role in obesity-associated OA and understanding of its mechanism are lacking. In addition, GAS6 has been described as a crucial bridging protein for macrophages to recognize and engulf apoptotic cells. Therefore, it was hypothesized that the high percentage of observed apoptotic cells might be due to ineffective macrophage efferocytosis caused by GAS6 suppression in OA. Primary BMDMs from ApoE^−/−^ mice and controls were fed CMFDA-FITC–labeled apoptotic cells (ACs) to determine the potency of efferocytosis induced by macrophages. The clearance of these fluorescent cells was quantified using flow cytometry. As expected, macrophage efferocytosis was impaired in the obesity microenvironment with reduced capacity for clearing fluorescent ACs in BMDMs from ApoE^−/−^ mice compared to the controls (Fig. 4E and F). Nevertheless, GAS6 had no obvious effect on the polarization of macrophages (Fig. 4G). Moreover, stimulation with GAS6 enhanced the phagocytotic activity of RAW264.7 cells, while inhibition of the Axl receptor by R428 diminished this effect (Fig. 4 H–J). Adding recombinant factor GAS6 significantly restored the up-regulation of inflammatory factors IL-1β, IL-6, and TNF-α induced by LPS in RAW264.7 cells (Supplementary Fig. S2C). These data indicate that M1 macrophage-associated reduction of GAS6 in obesity-associated OA mice promotes the accumulation of apoptotic cells by decreasing macrophage efferocytosis.

**Figure 4.**
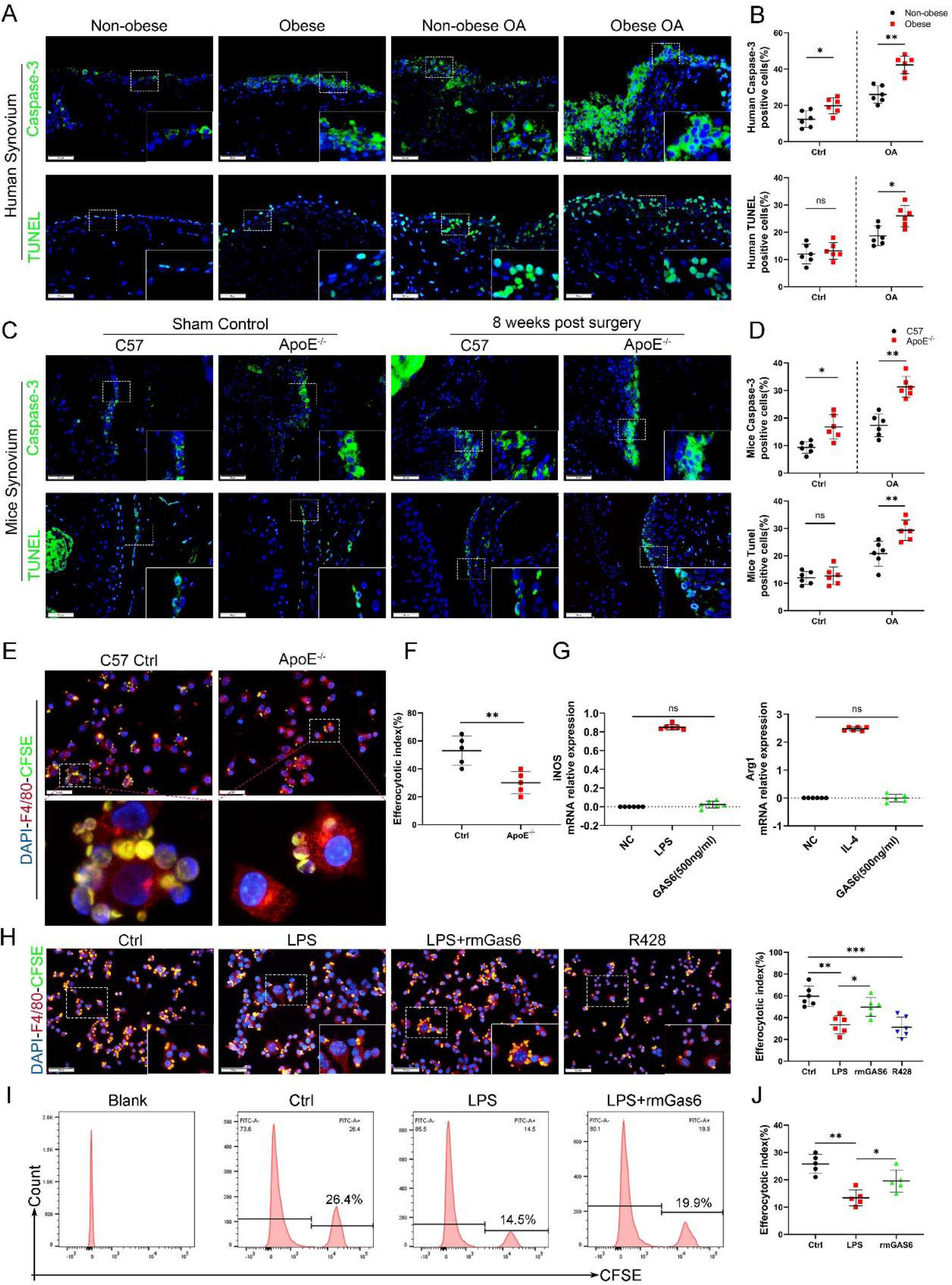
Accumulation of apoptotic cells in OA and impaired phagocytic ability of M1-polarized macrophages. (A) Immunofluorescence staining for caspase-3 (top) and TUNEL (lower) in normal and OA synovial tissue from non-obese and obese patients. Scale bar: 50 μm. Quantification of caspase-3- or TUNEL-positive cells as a proportion of total lining cell population in (A), n=6 per group. (C) Immunofluorescence staining for caspase-3 (top) and TUNEL (lower) in controls and DMM synovial tissue from C57 and ApoE−/− mice. Scale bar: 50 μm. (D) Quantification of caspase-3- or TUNEL-positive cells as a proportion of lining cell population in (C), n=6 per group. (E) Immunofluorescence staining for F4/80 (red) in BMDMs extracted from ApoE−/− and C57BL/6 mice. CFSE (green) in apoptotic thymocytes of C57BL/6 mice after 2-h phagocytosis. (F) Quantification of positive BMDMs engulfing apoptotic thymocytes as a proportion of total F4/80-positive cells, n=5 per group. (G) mRNA expression levels of iNOS or Arg1 after LPS, rmGAS6, or IL-4 stimulation for 24 h. (H) Immunofluorescence staining for F4/80 (red) in RAW264.7 cells and CFSE (green) in apoptotic thymocytes after phagocytosis for 2 h. Scale bar: 50 μm. Quantification of positive RAW264.7 cells engulfing apoptotic thymocytes as a proportion of total F4/80-positive cells, n=5 per group. (I) Flow cytometry analysis of CFSE-positive cells in total macrophages is shown as fluorescence-intensity distribution plots. (J) Efferocytotic index was calculated as percentage of CFSE-positive cells divided by percentage of total cells, n=6 per group. *P<0.05, **P<0.01, ***P<0.001, and NS=not significant. One-way analysis of variance (ANOVA) was performed. Data are shown as mean ± SD.

### 5. Suppression of GAS6/Axl axis promotes synovial hyperplasia, synovitis, and obesity-associated OA development

To further investigate the role of GAS6/Axl signaling in the development of OA *in vivo*, an intra-articular injection intervention was performed in OA mice. As a result, the degree of synovial inflammation and cartilage degeneration in C57BL/6 mice was far lower than that in ApoE^−/−^ mice, with a lower level of GAS6 expression in macrophages (Fig. 2A–C). Therefore, GAS6 recombinant factor was injected into ApoE^−/−^ obese OA mice to protect against synovial hyperplasia and cartilage damage induced by GAS6/Axl pathway suppression. On the other hand, the inhibitor R428 was injected into the joint cavity of C57BL/6 OA mice to stimulate OA progression. Interestingly, intra-articular injection of GAS6 significantly delayed synovial inflammation and cartilage destruction compared to vehicle-treated obese OA mice, manifested as lower synovitis and OARSI scores. In contrast, inhibition of Axl by R428 in C57BL/6 OA mice promoted synovial hyperplasia and cartilage destruction and enhanced synovitis and OARSI scores (Fig. 5A and B). Moreover, intra-articular injection of GAS6 recombinant factor decreased the number of apoptotic cells stimulated by synovitis in synovial tissues (Fig. 5C and D). Therefore, these data indicated that the GAS6/Axl axis might alleviate synovial hyperplasia and protect against obesity-associated OA development.

**Figure 5.**
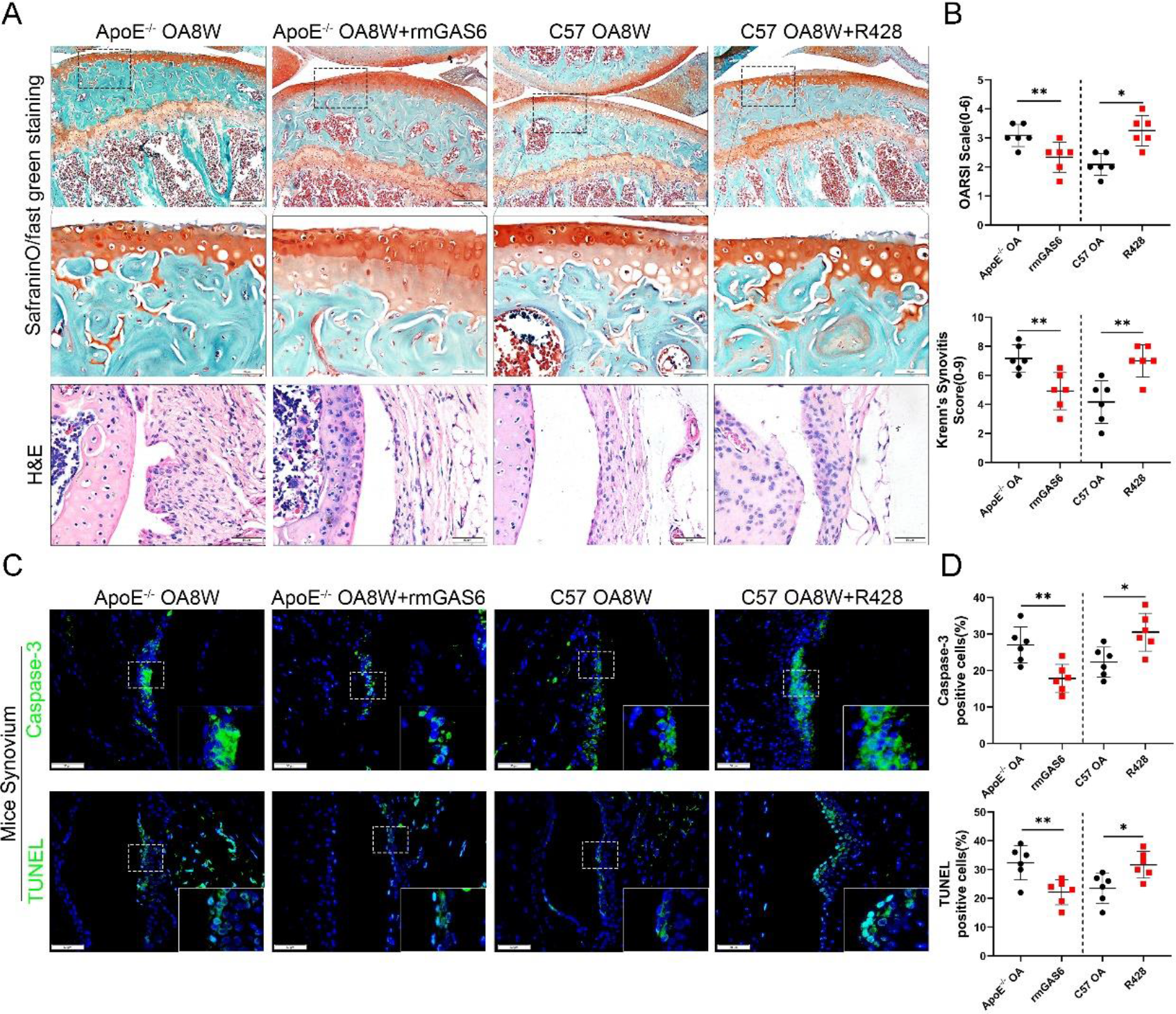
GAS6 restored OA cartilage loss and decreased apoptotic cell accumulation. (A) Safranin O and Fast Green staining (top and middle) of knee cartilage, H&E staining of synovial tissues from DMM mice and DMM mice treated with R428, and ApoE^−/−^ mice treated with recombinant mouse (rmGAS6) eight weeks after surgery. Scale bar: 200 μm, 50 μm. (B) Quantitative analysis of Osteoarthritis Research Society International (OARSI) scale and synovitis score in (A), n=6 per group. (C) Immunofluorescence staining of caspase-3 or TUNEL in synovial tissue from DMM mice, DMM mice treated with R428, and ApoE^−/−^ mice treated with recombinant mouse (rmGAS6) eight weeks after surgery. Scale bar: 50 μm. (D) Quantification of caspase-3- or TUNEL-positive cells as a proportion of lining cell population in (C), n=6 per group. *P<0.05, **P<0.01, ***P<0.001, NS=not significant. One-way analysis of variance (ANOVA) was performed. Data are shown as mean ± SD.

### 6. Model for obesity-associated synovitis and OA development

Obesity stimulates synovial macrophage infiltration and M1 polarization, which suppress the secretion of GAS6. GAS6 binds to the Axl receptor on macrophages, while GAS6/Axl inhibition promotes the accumulation of apoptotic cells by decreasing macrophage efferocytosis to induce chondrocyte degradation and aggravate OA development (Fig. 6).

**Figure 6.**
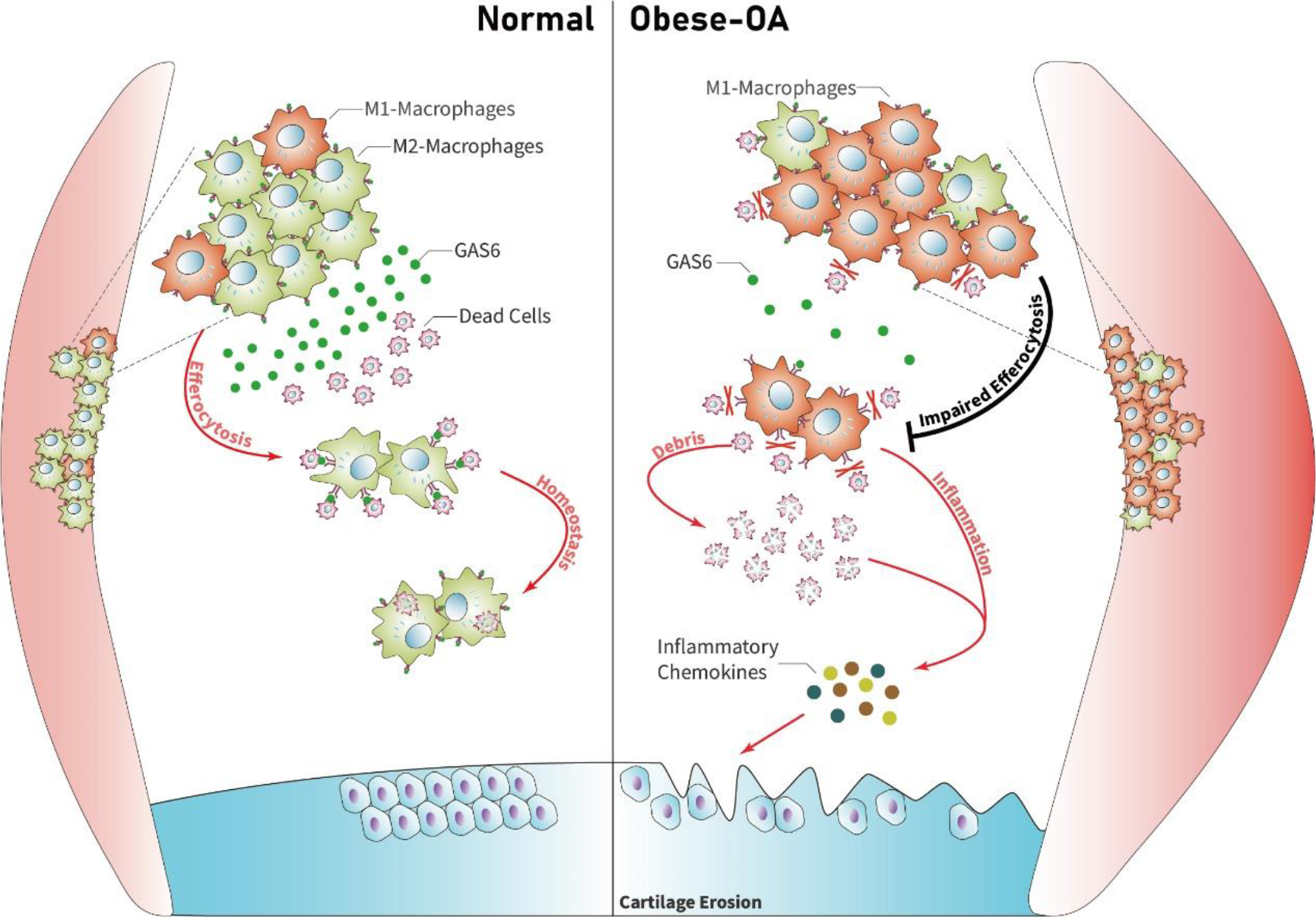
Model of GAS6 secreted by macrophages in modulating clearance of apoptotic cells and macrophage polarization during OA. Macrophage polarization induced by obesity decreased the secretion of GAS6 and impaired the phagocytosis of apoptotic cells. The accumulation of apoptotic cell debris lead to the persistence of local inflammation and synovial hyperplasia, which aggravates the pathological process of OA.

## Discussion

The present study revealed for the first time that M1-polarized macrophage infiltration in OA synovial tissue of obese patients is increased, accompanied by markedly down-regulated secretion of GAS6 and impaired macrophage-dependent efferocytosis for cleaning apoptotic cells. The intracellular contents released by accumulated apoptotic cells further trigger an immune response and lead to a release of inflammatory factors, such as TNF-α, IL-1β, and IL-6, which induce the dysfunction of chondrocyte homeostasis in obesity-associated OA. Therefore, blocking M1 macrophage polarization or intra-articular injection of GAS6 is a potential therapeutic strategy for obesity-associated OA.

Obesity has always been considered a significant risk factor in the progression of OA^27^, leading to more severe OA manifestations, including cartilage loss, subchondral bone sclerosis, and synovial inflammation.^13^ Researchers have accepted the role of synovial inflammation in the pathological progression of OA.^16,28^ However, the underlying mechanism of obesity-related inflammation in the development of synovitis in OA remains unclear. The present study revealed that obese patients and ApoE^−/−^ mice showed a more severe cartilage destruction with enhanced OARSI scores.^29^ Furthermore, the lining layer of synovial tissue was more prone to hyperplasia. The number of macrophages increased significantly during the pathological development of OA in obese patients, which manifested as enhanced synovitis scores and increased numbers of F4/80-positive cells.^30^ These results suggest that obesity plays an essential mediating role in OA development.

Recent studies have reported that the imbalance in M1/M2 macrophage polarization plays a vital role in developing OA inflammation.^19^ Classically, macrophages are divided into inflammatory M1 and anti-inflammatory M2 macrophages,^31^ though they can be interchanged or transformed into each other during various inflammatory reactions and thus function differently.^32^ Lumeng et al. have found that adipose tissue macrophages (ATMs) prefer to express TNF-α, iNOS, and other M1 macrophage markers, while ATMs in non-obese individuals highly express M2 macrophage markers.^33^ However, the effect of obesity on inducing macrophage polarization during OA development remains unclear. Many bursas and adipose tissue around the knee joint are typically characterized as the infrapatellar fat pad.^34^ Some researchers believe that the transformation of macrophages is triggered by lipids released by fat cells at the onset of obesity.^35^ Thus, we investigated whether macrophage abnormalities in obesity-related knee OA affect the synovial membrane in the knee joint.

We found that M1 macrophage infiltration increased in the OA synovial tissue of obese patients and ApoE^−/−^ mice compared to non-obese patients and C57BL/6 mice, accompanied by increased secretion of TNF-α, IL-1, and IL-6. These results suggest that obesity may be a crucial factor affecting the functional status of macrophages, and targeting the polarization of macrophages in obese patients may alleviate the synovitis in OA.

The balance of progression and regression that controls the state of local inflammation has recently been in the spotlight.^36,37^ The maintenance of efferocytosis dampens pro-inflammatory cytokine production and initiates inflammation resolution.^25,38^ Emerging studies have mentioned the effect of macrophage polarization on phagocytic ability. Yurdagul et al. have shown that M2 macrophages retain efferocytosis properties, which are crucial for resolving inflammation.^39^ Another study demonstrated that DEL-1 contributes to the resolution of inflammation by promoting apoptotic neutrophil efferocytosis through macrophages and the emergence of an M2 macrophage phenotype.^40^ Although efferocytosis has been widely studied in multiple disease models, its role in synovial inflammation and OA development has not been reported. Our *in vitro* experiments demonstrated that macrophages extracted from the bone marrow of obese mice had decreased phagocytic capacity, and the phagocytic ability of macrophages for apoptotic thymocytes was reduced after stimulation with LPS (500 ng/mL). These results showed that the proportion of apoptotic cells was significantly increased in the synovium of ApoE^−/−^ mice. This is partly due to the decreased phagocytic ability caused by the enhanced M1 macrophage polarization.

Intra-articular injection of GAS6 restored the phagocytic capacity of macrophages. It reduced the accumulation of local apoptotic cells and decreased the levels of TUNEL- and caspase-3-positive cells, preserving cartilage thickness and preventing the progression of obesity-associated OA. These findings suggest that targeting the efferocytosis of macrophages in local inflammation of obesity-associated synovitis may maintain the homeostasis of the cartilage cavity and alleviate the obesity-related OA.

GAS6 and its receptor Axl are known to regulate apoptotic-related inflammation and may be implicated in lupus pathogenesis.^41^ The remaining apoptotic cells are a source of autoantigens and can drive autoimmunity development.^42^ The expression of GAS6 in the hyperplasia synovial tissues from obesity-associated OA in the present study was down-regulated and accompanied by an increase in M1 macrophage polarization. Moreover, the *in vitro* experiments demonstrated a decreased secretion of GAS6 protein and impaired phagocytic ability in macrophages after LPS stimulation. However, the proportion of macrophages that engulfed apoptotic cells was increased after incubation with recombinant GAS6 protein, while the phagocytic ability was significantly down-regulated after blocking the Axl receptor by adding R428. Exogenous cultured rmGAS6 with macrophages after stimulation of LPS can also decrease the levels of inflammatory cytokines, such as IL-1, IL-6, and TNF-ɑ. Nevertheless, there was no significant difference in the expression of its specific receptor Axl, which may be explained by the fact that GAS6 has three receptors (Axl, Mer, and Tyro3) with different affinities. These findings suggest that GAS6 may relieve local synovial inflammation by restoring the phagocytic ability of macrophages for apoptotic cells and decrease the induction of inflammatory chemokines, which alleviate the pathological progression of OA.

To conclude, the present study found that obese OA patients and ApoE^−/−^ obese mice showed a more pronounced synovitis and enhanced macrophage infiltration in synovial tissue, accompanied by dominant M1 macrophage polarization. Obese OA mice had more severe cartilage destruction than OA mice in the control group. Enhanced M1-polarized macrophages in obese synovium decreased GAS6 secretion, impairing efferocytosis for synovial apoptotic cells and causing synovial hyperplasia and obesity-associated OA development. Therefore, these findings reveal that targeting GAS6-mediated macrophage polarization and phagocytosis in obese patients with OA may be a potential therapeutic strategy.

## Materials and methods

### Human synovial tissue

Normal human synovial tissue and synovial fluid samples were obtained from patients requiring arthroscopic treatment of acute anterior cruciate ligament rupture or meniscus injury (n = 6, age 34 ± 8.15 years, three males, three females). Other joint diseases were excluded from the study. Synovial OA tissues were obtained from patients undergoing total knee arthroplasty (n = 6, age 64 ± 3.58 years, two males, four females). Informed consent was obtained from all recruited patients and was identified by the ethics committee of the Third Affiliated Hospital of Southern Medical University.

### Destabilization of medial meniscus (DMM) animal model

Ten-week-old male C57BL/6 mice (body weight 23 ± 2 g) and ApoE-deficient (ApoE^−/−^) male mice (body weight 33 ± 2 g) were purchased from the Experimental Animal Center of Guangdong Province, China. All animals were housed in cages without pathogens at a temperature of 24 ± 5°C and with a relative humidity of 40%. C57BL/6 mice were fed a standard diet, and ApoE^−/−^ mice were fed a high-fat diet. The feed specifications are shown in Table 2. The protocol was approved by the Southern Medical University Animal Care and Use Review Board.

In the DMM-OA model, mice were anesthetized by intraperitoneal injection of 5% chloral hydrate and the skin was cut along the medial collateral ligament. The joint capsule was cut open and the femoral condyle was exposed. The connection between the medial meniscus and the tibial plateau was cut to release the medial meniscus. The joint capsule and skin were sutured after the operation.

### Animal treatment and specimen preparation

After the surgery, 50 ng/g of recombinant mouse GAS6 (rmGAS6, Sino Biological, China, #58026-M08H) was administered into the articular once per week. The right legs were harvested four or eight weeks post-surgery (n = 6 in each group). Knee joints from mice in different experimental groups were fixed in 4% paraformaldehyde for 48 h and decalcified for 21 days. The specimens were embedded in paraffin, and 4-μm serial sections were cut from the sagittal portion through the inner side of the knee. The Southern Medical University Animal Care and Use Committee approved all procedures involving mice.

### Histology and immunohistochemical (IHC)/immunofluorescence (IF) staining

Histology sections were stained with Safranin O-fast green/hematoxylin and eosin (H&E) for morphological analysis. IHC and IF staining was performed on the 4-μm-thick tissue sections. Slides were deparaffinized, rehydrated, and washed in phosphate-buffered saline (PBS) three times for 5 min each time. Antigen retrieval was performed by soaking slides in citric acid overnight in a 60°C water bath. After washing three times in PBS, slides were quenched in 3% hydrogen peroxide for 10 min at room temperature and washed with PBS three more times. Then, slides were blocked with 10% normal bovine serum (Solarbio, Beijing, China) for 1 h at room temperature (IHC staining). Slides were then incubated with primary antibodies at 4°C overnight. A secondary antibody for IHC or fluorescent secondary antibody for IF was applied for 1 h at room temperature. Then, IHC slides were stained with diaminobenzidine and hematoxylin, dehydrated, and mounted. IF slides were processed with 4, 6-diamidino-2-phenylindole (DAPI, Thermo Fisher Scientific, Waltham, MA, USA) staining solution and mounted with cover glass. Antibodies used for IHC/IF staining were as follows: rabbit anti-caspase-3 (Proteintech, 1:200, #19677-1-AP), rabbit anti-GAS6 (Abclone, 1:100, #A8545), rabbit anti-Axl (Abclone, 1:100, #A20548), rabbit anti-MMP13 (Proteintech, 1:400, 18165-1-AP), rabbit anti-aggrecan (Proteintech, 1:400, 13880-1-AP), mouse anti-F4/80 (Santa Cruz Biotechnology, 1:100, sc377009), mouse anti-iNOS (Santa Cruz Biotechnology, 1:100, sc-7271), mouse anti-CD206 (Proteintech, 1:100, 18704-1-AP), species-matched horseradish peroxidase-conjugated secondary antibodies (Jackson Immuno Research Laboratories), and species-matched Alexa-488 or 594-labeled secondary antibody (Life Technologies, Carlsbad, CA, USA).

### Cartilage and synovium structure grading

Histology sections of the knee joints were graded based on the Osteoarthritis Research Society International (OARSI) scoring system developed by Glasson et al.^1^ by two observers blinded to the experimental conditions. Generally, sections were assigned a grade of 0–6: 0, normal cartilage; 0.5, slight loss of Safranin O staining without structural changes; 1, small fibrillations without loss of cartilage; 2, vertical clefts down to the layer below the superficial layer; 3–6, vertical clefts or erosion to the calcified cartilage affecting <25% (grade 3), 25–50% (grade 4), 50–75% (grade 5), and >75% (grade 6) of the articular surface. Synovitis severity was estimated based on the synovial lining cell layer enlargement, resident cell density, and inflammatory infiltration. A nine-point scale was used, where low scores indicated moderate synovitis and high scores represented severe synovitis.^2^

### TUNEL

The TUNEL assay was performed according to the manufacturer’s instructions (TUNEL Apoptosis Detection Kit (Alexa Fluor 488), Yeasen, #40307ES20) to detect cell death in the synovial membrane. The assay used the green channel at 488 nm. DAPI was applied as a nuclear counterstain in the blue channel at 461 nm. Images were taken with an *Olympus BX43* fluorescent microscope and *Olympus DP73* digital camera at 400× magnification with cellSens software (Olympus). Exposure settings were adjusted to minimize oversaturation.

### Apoptosis induction and thymocyte immunofluorescence staining

Murine thymocytes were isolated from C57BL/6 mice and then stimulated with 25 μmol/L dexamethasones for 3 h to induce apoptosis, followed by washing twice and resuspension with phagocyte culture medium to a concentration of 1 × 10^7^ cells/mL. Then, thymocytes were labeled with carboxyfluorescein succinimidyl ester (CFSE) dye (Topscience, #T6802) following the manufacturer’s instructions.

### Efferocytosis assay

An efferocytosis assay was performed as previously described.^3^ Briefly, RAW264.7 cells (ATCC, Manassas, VA, USA) were plated in a six-well plate (1 × 10^6^ cells/well) with Dulbecco’s Modified Eagle Medium containing 10% fetal bovine serum and cultured overnight. The cells were then treated with LPS (Invitrogen, San Diego, CA, USA), rmGAS6, or R428 (Topscience, #CAS 1037624-75-1) for 24 or 48 h. CFSE-labeled apoptotic thymocytes were then added at a ratio of 10:1 and incubated for an additional 120 min. The cells were extensively washed three times with PBS to remove unengulfed thymocytes. The ability of macrophages to engulf apoptotic thymocytes was quantified by flow cytometry or visualized using immunofluorescence microscopy.

The efferocytotic index was calculated using the following formula: (number of macrophages containing apoptotic bodies) / (total macrophages) × 100% and then normalized using the control group as 100%.

### Real-time polymerase chain reaction (RT-PCR)

Total RNA was isolated from RAW264.7 cells and ground cartilage from human tibial plateaus using TRIzol reagent (Takara Bio Inc., Shiga, Japan). For mRNA quantification, 1 mg of total RNA was purified with gDNA remover and reverse transcribed using 5× HiScript II qRT SuperMix II (Vazyme Biotech, Nanjing, China). Each PCR reaction consisted of 10 μL of 2× ChamQ SYBR qPCR Master Mix (Vazyme), 10 μM of forward and reverse primers, and 500 ng of cDNA. For miRNA quantification, 1 mg of total RNA was purified with gDNA wiper mix and then reverse transcribed using Hiscript II Enzyme Mix, 10× RT Mix, and specific stem-loop primers. Template DNA was mixed with 2× miRNA Universal SYBR qPCR Master Mix, specific primers, and mQ primer R (Vazyme). All reactions were run in triplicate.

Mouse primer sequences are listed below:

GAS6

Forward: 5′- CCGCGCCTACCAAGTCTTC-3’

Reverse: 5′-CGGGGTCGTTCTCGAACAC-3’

GAPDH

Forward 5′-AAATGGTGAAGGTCGGTGTGAAC-3’

Reverse 5′-CAACAATCTCCACTTTGCCACTG-3’

IL-1β

Forward 5′-GCAACTGTTCCTGAACTCAACT-3’

Reverse 5′-ATCTTTTGGGGTCCGTCAACT-3’

IL-6

Forward 5′-ACAACCACGGCCTTCCCTACTT-3’

Reverse 5′-CAGGATTTCCCAGCGAACATGTG-3’

TNF-α

Forward 5′CCTCCCTCTCATCAGTTCTA-3’

Reverse 5′-ACTTGGTTTGCTACGAC-3’

### Enzyme-linked immunosorbent assay (ELISA)

Human synovial fluid samples were collected as described above. All samples were spun down at 4500 × *g* for 15 min. Human GAS6 Quantikine Kit (R&D Systems) was used to measure the concentration of GAS6 in the synovial fluid.

### Statistical analyses

Data were represented as the mean ± SD. An unpaired Student’s t-test was performed for experiments comparing two groups of data. A one-way analysis of variance (ANOVA) was performed for data involving multiple groups, followed by Tukey’s post-hoc test. P-values of <0.05 were considered statistically significant.

## Acknowledgments

This work was supported by grants from the National Natural Science Foundation of China (Grant numbers: 81902229 and 81871745).

## Competing Interests

The authors have declared that no conflict of interest exists.

## Author contributions

HZ and XB conceived the ideas for experimental designs, analysed data and wrote the manuscript. ZY, WQ and LL conducted the majority of the experiments and helped with manuscript preparation. AL, YS, and HZ performed immunohistochemistry and immunofluorescence and confocal imaging. HZ and HP conducted cell cultures and western blot experiments. XG and JY collected human tissue samples. HZ and DC developed the concept, supervised the project and conceived the experiments.

## Data availability statement

All data generated or analyzed during this study are included in this submitted article and its additional files.

---

**Supplementary Figure S1.**
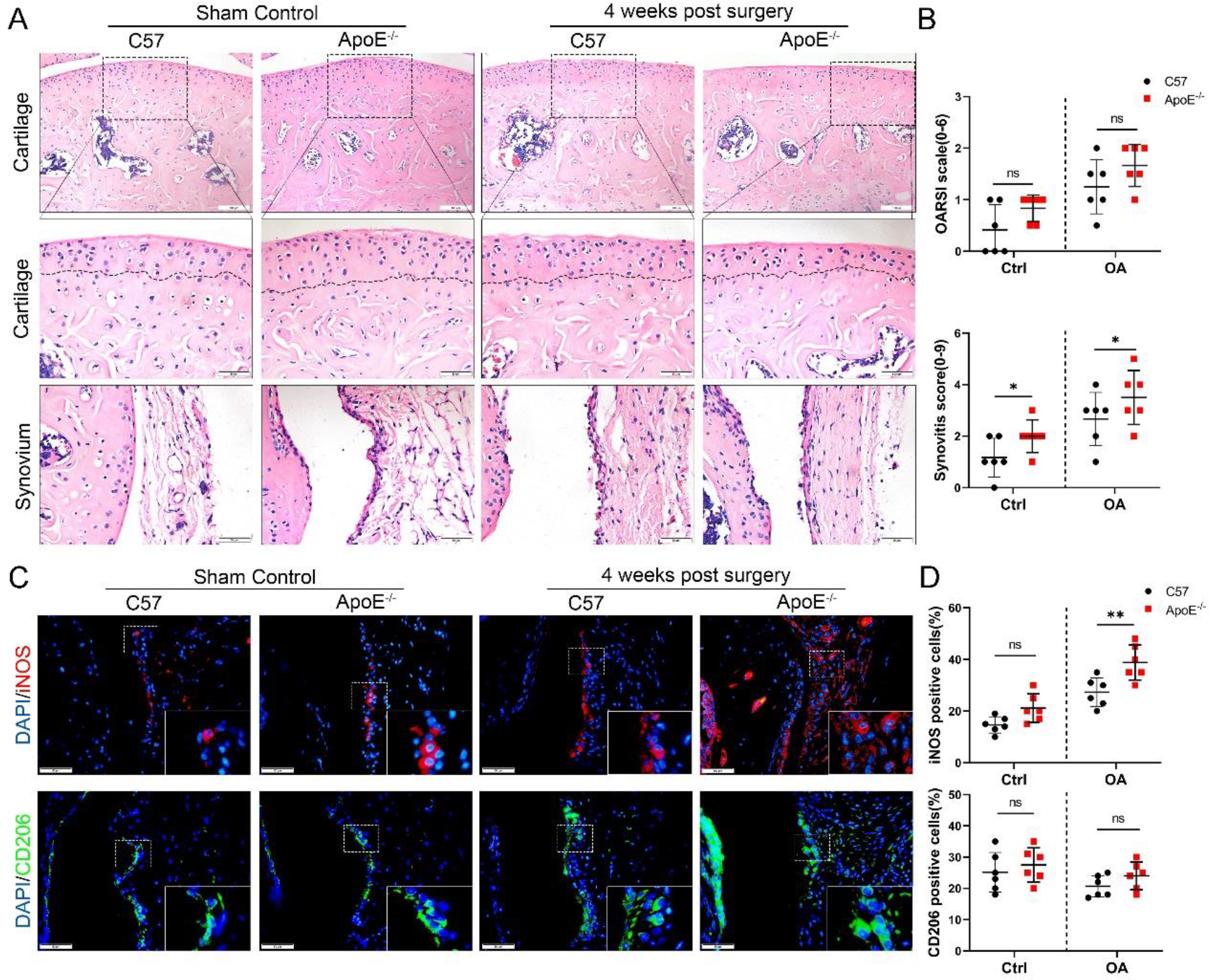
Cartilage loss, synovial hyperplasia, and macrophage polarization in ApoE^−/−^ OA. (A) H&E staining of cartilage and synovial tissue in controls and DMM from normal and ApoE^−/−^ mice four weeks after surgery. Scale bar: 200 μm, 50 μm. (B) Quantitative analysis of Osteoarthritis Research Society International (OARSI) scale and synovitis score described in (A). (C) Immunofluorescence of iNOS and CD206 in controls and DMM synovial tissues from normal and ApoE^−/−^ mice four weeks after surgery. Scale bar: 50 μm. (D) Quantification of iNOS- and CD206-positive cells as a proportion of lining cells in (C), n=6 per group. *P<0.05, **P<0.01, ***P<0.001, NS=not significant. One-way analysis of variance (ANOVA) was performed. Data are shown as mean ± SD.

**Supplementary Figure S2.**
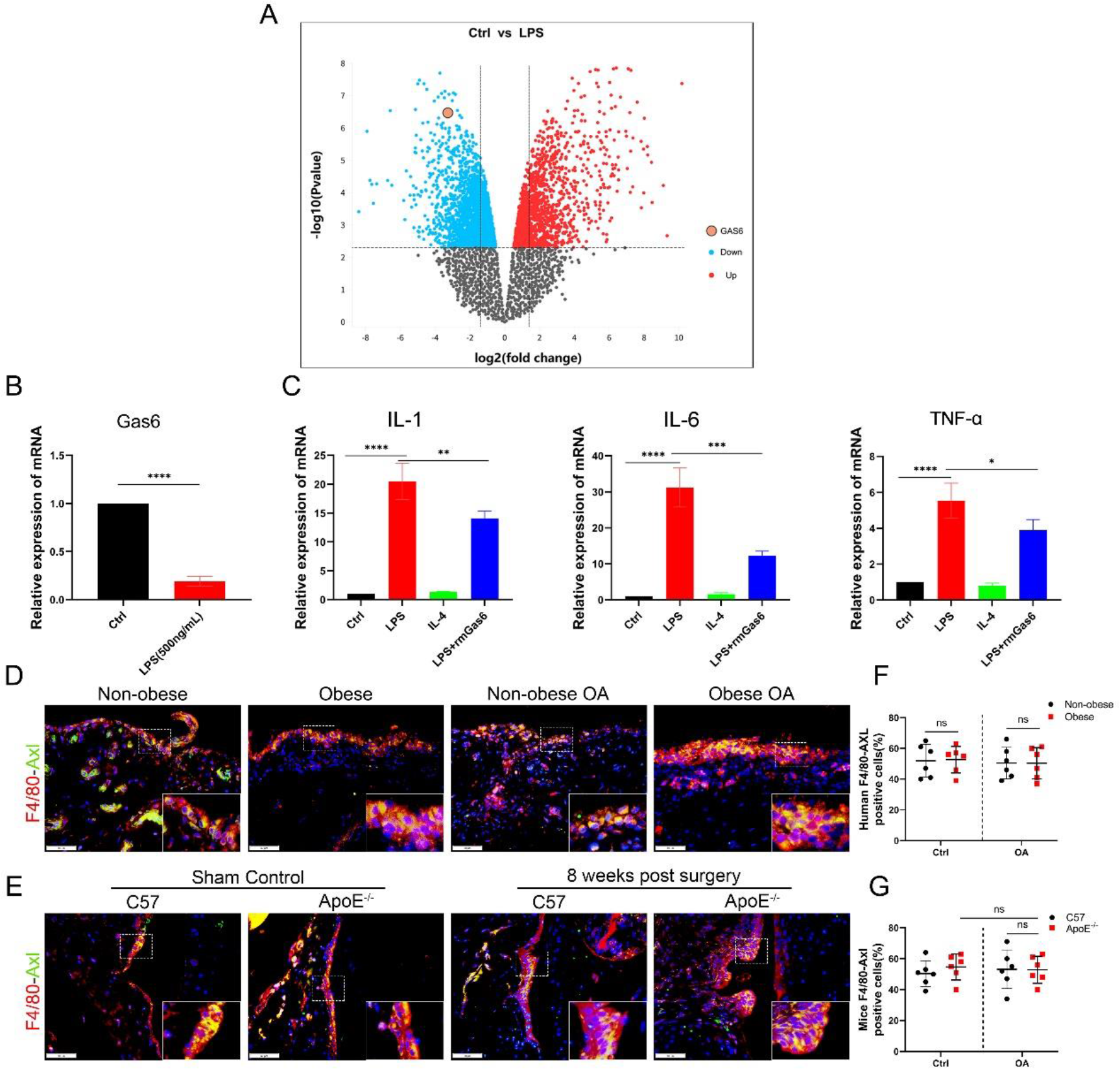
(A) Differentially-expressed mRNA in bone marrow-derived macrophages from normal controls or lipopolysaccharide treatment based on GSE53986. (B) Relative mRNA expression level of GAS6 in LPS-treated RAW264.7 cells, n=3 per group. (C) Relative mRNA expression level of IL-1β, IL-6, and TNF-α in LPS and rmGAS6-treated RAW264.7 cells, n=3 per group. (D) Immunofluorescence of F4/80 (red) and AXL (green) in synovial tissue from non-obese human, non-obese OA patients, obese human, and obese OA patients. Scale bar: 50 μm. (E) Immunofluorescence staining of F4/80 (red) and AXL (green) in synovial tissue in controls and DMM cartilage from C57BL/6 and ApoE^−/−^ mice. Scale bar: 50 μm; (F and G) Quantification of F4/80-AXL-positive macrophages (yellow) as a proportion of total F4/80-positive cells in (D and E), n=6 per group. *P<0.05, **P<0.01, NS=not significant. One-way analysis of variance (ANOVA) was performed. Data are shown as mean ± SD.

## Figures (Supplementary figures and tables)

**Table 1.**
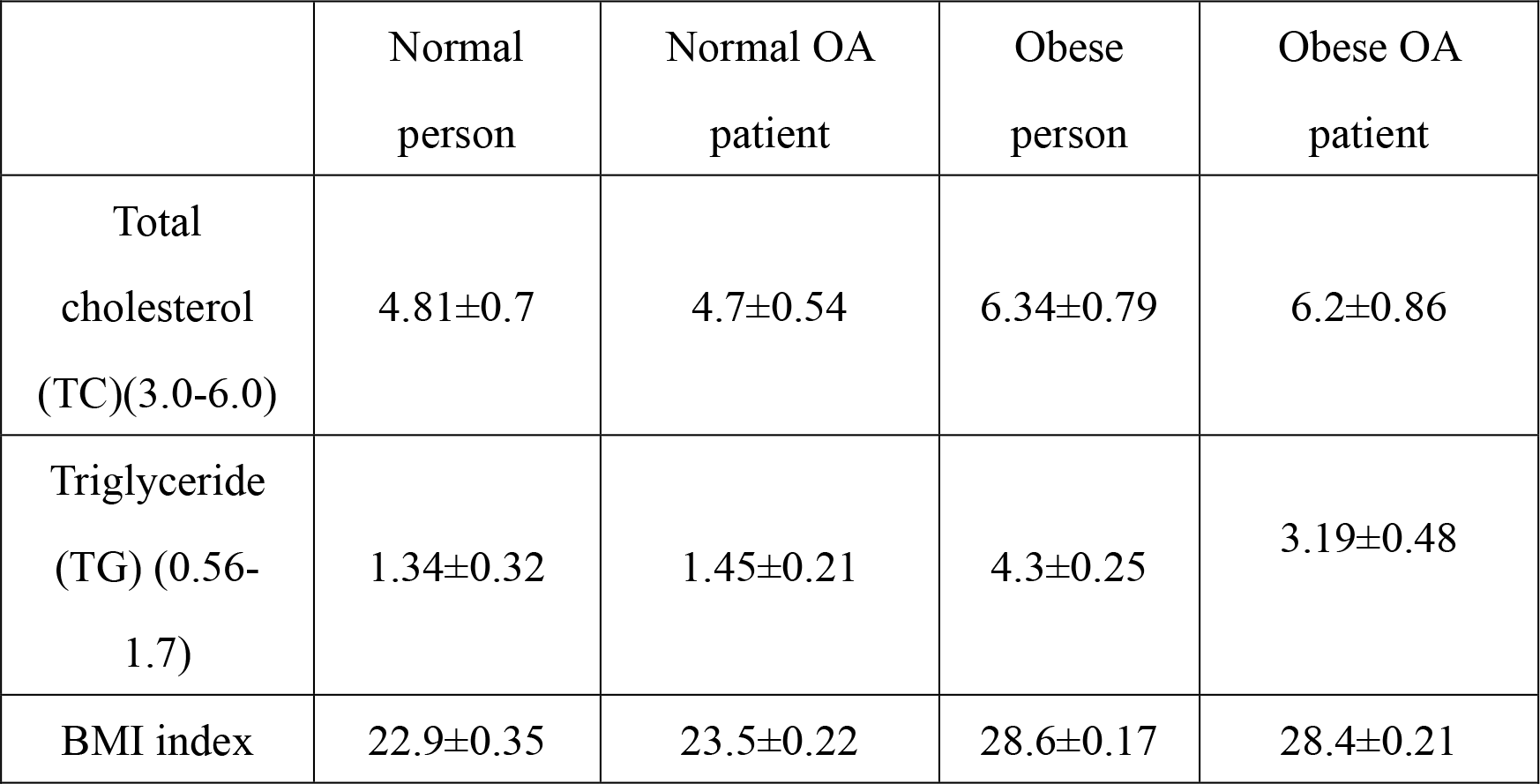
Blood lipids in obesity-related patients and general patients

**Table 2.**
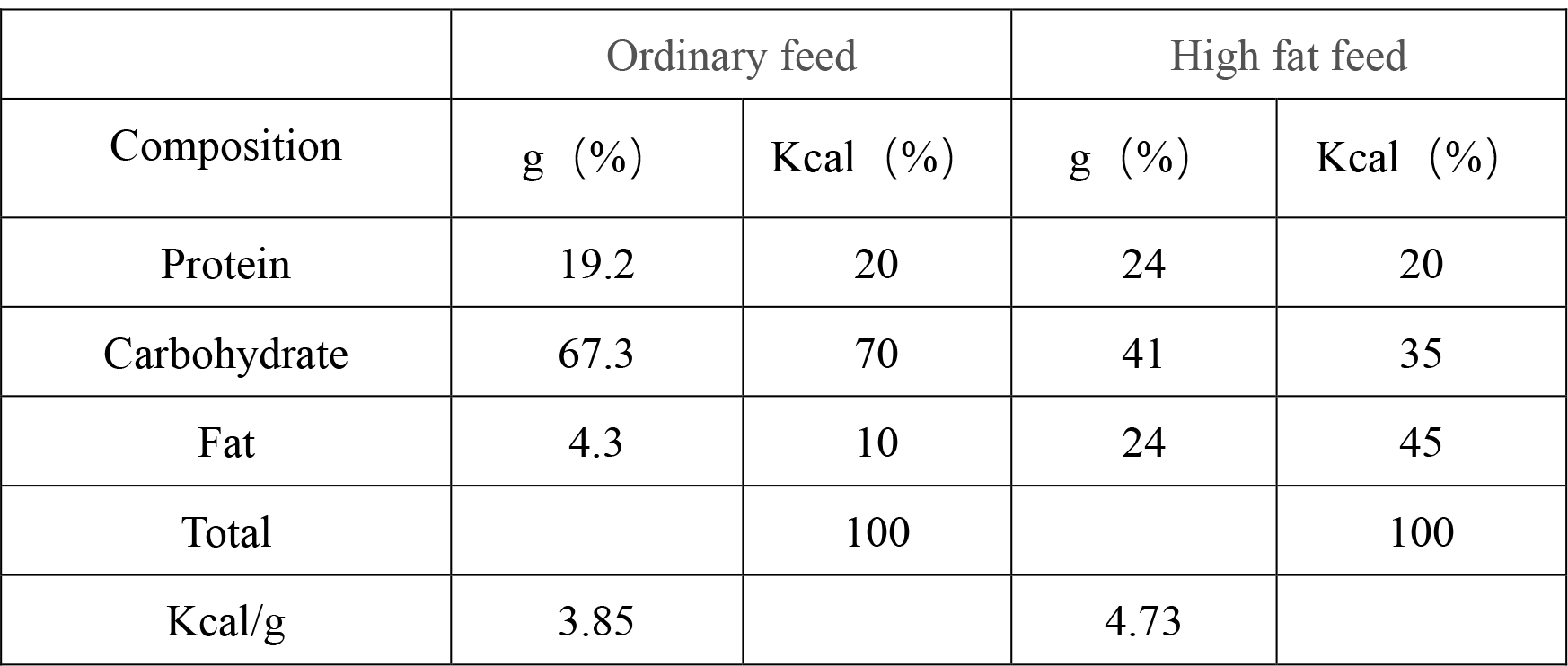
Comparison of specifications and energy of ordinary feed and high fat feed

|  | Ordinary feed |  | High fat feed |  |
| --- | --- | --- | --- | --- |
| Composition | g (%) | Kcal (%) | g (%) | Kcal (%) |
| Protein | 19.2 | 20 | 24 | 20 |
| Carbohydrate | 67.3 | 70 | 41 | 35 |
| Fat | 4.3 | 10 | 24 | 45 |
| Total |  | 100 |  | 100 |
| Kcal/g | 3.85 |  | 4.73 |  |

**Table 3.** Lipid status of APOE^−/−^ obese mice and C57BL/6 mice

|  | APOE<sup>-/-</sup> mice | C57BL/6 mice |
| --- | --- | --- |
| Total cholesterol (TC)<br>(3.0-6.0) | 17.82±3.3 | 2.77±0.62 |
| Triglyceride (TG)<br>(0.56-1.7) | 2.56±0.43 | 0.92±0.31 |

**Table 4.**
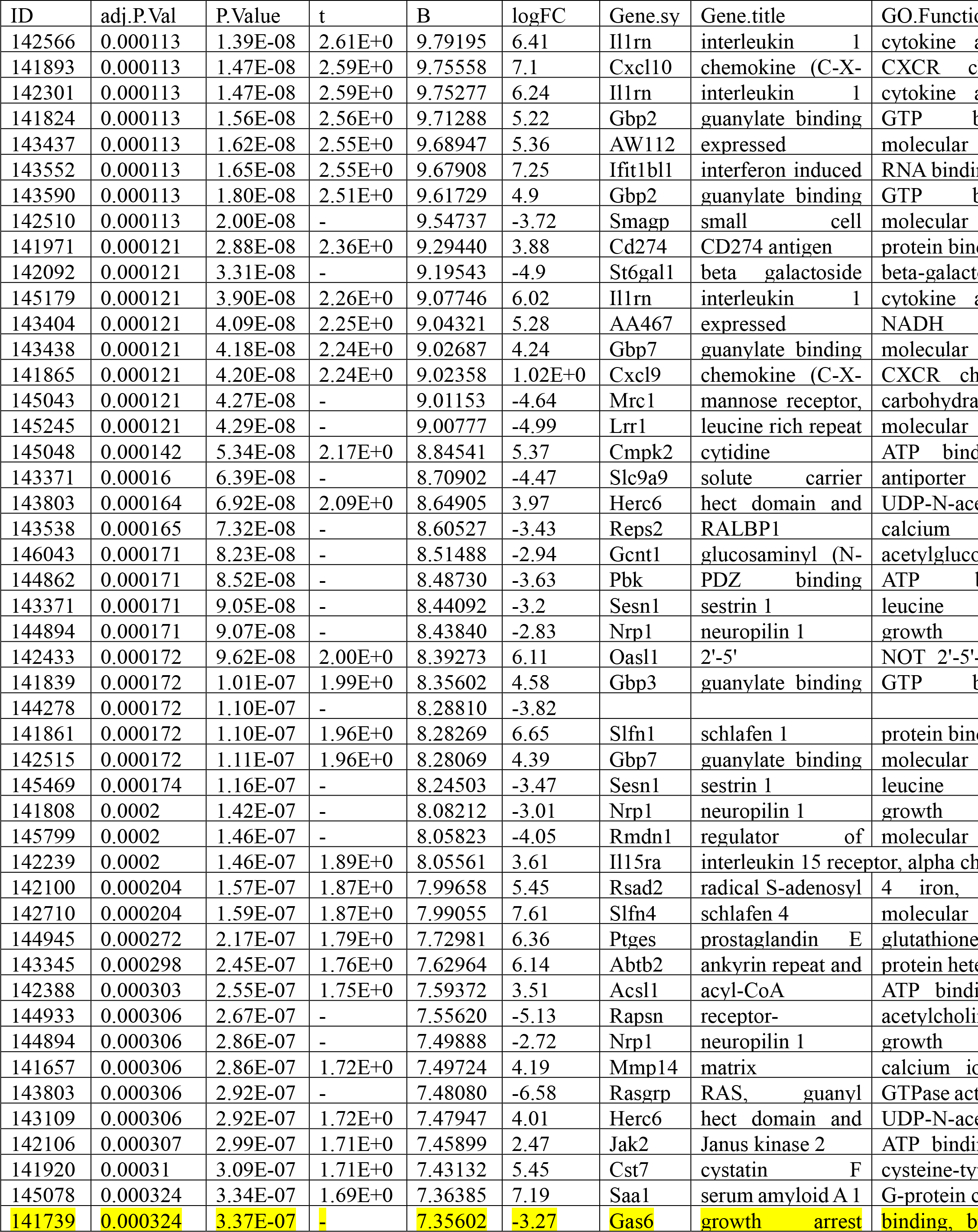
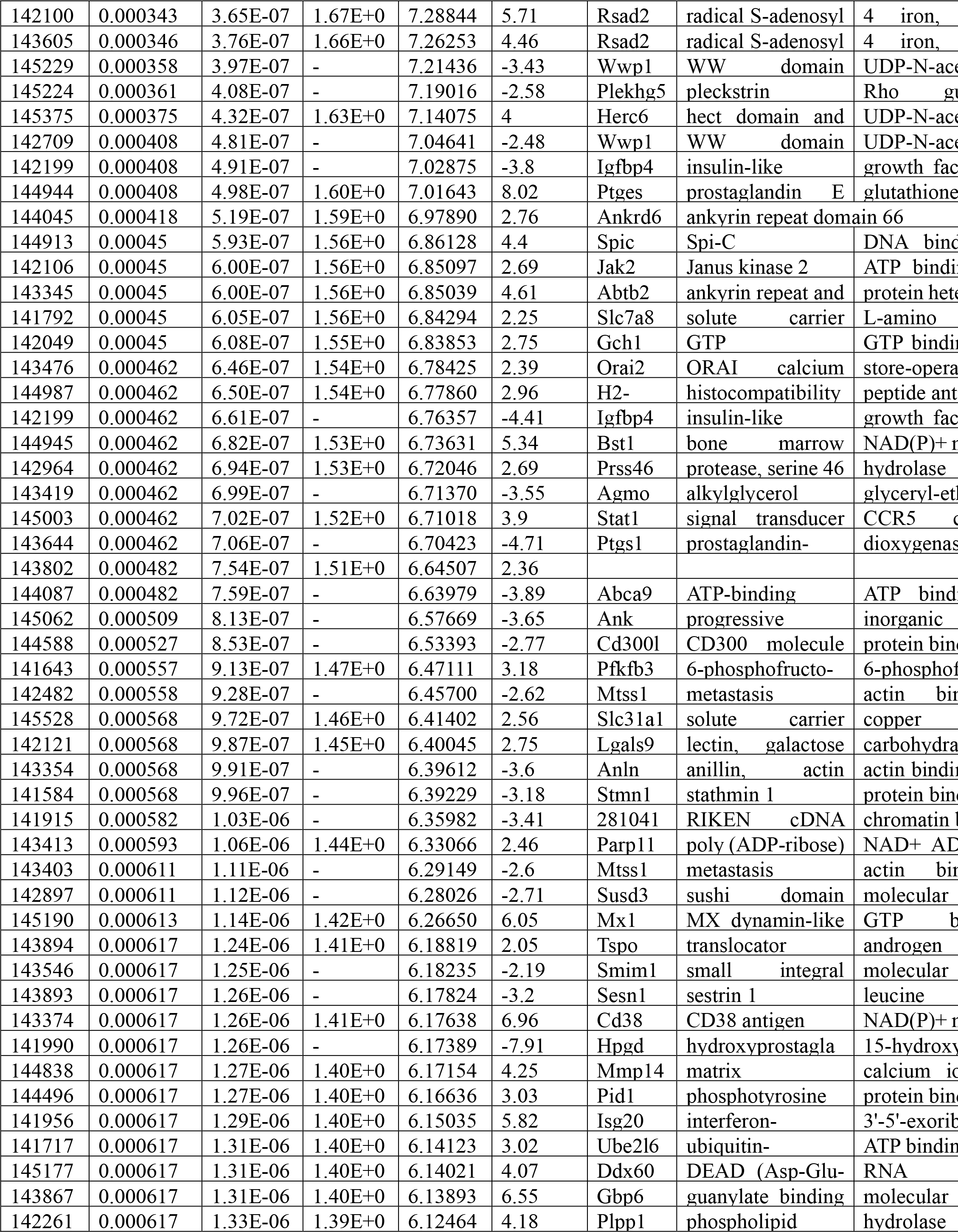
Results of GSE53986

## Notes

### Competing Interest Statement

The authors have declared no competing interest.

## References

1. Hunter, D. J. & Bierma-Zeinstra, S. Osteoarthritis. The Lancet 393, 1745–1759 (2019).

2. Martel-Pelletier, J. et al. Osteoarthritis. Nat. Rev. Dis. Primer 2, 16072 (2016).

3. Teichtahl, A. J. et al. Weight change and change in tibial cartilage volume and symptoms in obese adults. Ann. Rheum. Dis. 74, 1024–1029 (2015).

4. Clement, N. D. & Deehan, D. J. Overweight and Obese Patients Require Total Hip and Total Knee Arthroplasty at a Younger Age. J. Orthop. Res. Off. Publ. Orthop. Res. Soc. 38, 348–355 (2020).

5. Franklin, J., Ingvarsson, T., Englund, M. & Lohmander, L. S. Sex differences in the association between body mass index and total hip or knee joint replacement resulting from osteoarthritis. Ann. Rheum. Dis. 68, 536–540 (2009).

6. Caballero, B. Humans against Obesity: Who Will Win? Adv. Nutr. Bethesda Md 10, S4–S9 (2019).

7. Trends in adult body-mass index in 200 countries from 1975 to 2014: a pooled analysis of 1698 population-based measurement studies with 19·2 million participants. Lancet Lond. Engl. 387, 1377–1396 (2016).

8. Yusuf, E. et al. Association between weight or body mass index and hand osteoarthritis: a systematic review. Ann. Rheum. Dis. 69, 761–765 (2010).

9. Visser, A. W. et al. Adiposity and hand osteoarthritis: the Netherlands Epidemiology of Obesity study. Arthritis Res. Ther. 16, R19 (2014).

10. Visser, A. et al. Adiposity and hand osteoarthritis: the Netherlands Epidemiology of Obesity study. Arthritis Res. Ther. 16, R19 (2014).

11. Pottie, P. et al. Obesity and osteoarthritis: more complex than predicted! Ann.Rheum. Dis. 65, 1403–1405 (2006).

12. Neumann, E., Junker, S., Schett, G., Frommer, K. & Müller-Ladner, U. Adipokines in bone disease. Nat. Rev. Rheumatol. 12, 296–302 (2016).

13. Wang, T. & He, C. Pro-inflammatory cytokines: The link between obesity and osteoarthritis. Cytokine Growth Factor Rev. 44, 38–50 (2018).

14. Xie, C. & Chen, Q. Adipokines: New Therapeutic Target for Osteoarthritis? Curr. Rheumatol. Rep. 21, 71 (2019).

15. Sellam, J. & Berenbaum, F. The role of synovitis in pathophysiology and clinical symptoms of osteoarthritis. Nat. Rev. Rheumatol. 6, 625–635 (2010).

16. Mathiessen, A. & Conaghan, P. G. Synovitis in osteoarthritis: current understanding with therapeutic implications. Arthritis Res. Ther. 19, 18 (2017).

17. Koski, J. M. et al. Power Doppler ultrasonography and synovitis: correlating ultrasound imaging with histopathological findings and evaluating the performance of ultrasound equipments. Ann. Rheum. Dis. 65, 1590–1595 (2006).

18. Daghestani, H. N., Pieper, C. F. & Kraus, V. B. Soluble macrophage biomarkers indicate inflammatory phenotypes in patients with knee osteoarthritis. Arthritis Rheumatol. Hoboken NJ 67, 956–965 (2015).

19. Zhang, H. et al. Synovial macrophage M1 polarisation exacerbates experimental osteoarthritis partially through R-spondin-2. Ann. Rheum. Dis. 77, 1524–1534 (2018).

20. Sun, Y., Zuo, Z. & Kuang, Y. An Emerging Target in the Battle against Osteoarthritis: Macrophage Polarization. Int. J. Mol. Sci. 21, (2020).

21. Collins, K. H. et al. Adipose tissue is a critical regulator of osteoarthritis. Proc. Natl. Acad. Sci. U. S. A. 118, (2021).

22. Bellan, M. et al. Gas6/TAM System: A Key Modulator of the Interplay between Inflammation and Fibrosis. Int. J. Mol. Sci. 20, 5070 (2019).

23. Doran, A. C., Yurdagul, A. & Tabas, I. Efferocytosis in health and disease. Nat. Rev. Immunol. 20, 254–267 (2020).

24. Nepal, S. et al. STAT6 induces expression of Gas6 in macrophages to clear apoptotic neutrophils and resolve inflammation. Proc. Natl. Acad. Sci. 116, 16513–16518 (2019).

25. Boada-Romero, E., Martinez, J., Heckmann, B. L. & Green, D. R. The clearance of dead cells by efferocytosis. Nat. Rev. Mol. Cell Biol. 21, 398–414 (2020).

26. Noubade, R. et al. NRROS negatively regulates reactive oxygen species during host defence and autoimmunity. Nature 509, 235–239 (2014).

27. Kulkarni, K., Karssiens, T., Kumar, V. & Pandit, H. Obesity and osteoarthritis. Maturitas 89, 22–28 (2016).

28. Felson, D. T. et al. Synovitis and the risk of knee osteoarthritis: the MOST Study. Osteoarthritis Cartilage 24, 458–464 (2016).

29. Glasson, S. S., Chambers, M. G., Van Den Berg, W. B. & Little, C. B. The OARSI histopathology initiative – recommendations for histological assessments of osteoarthritis in the mouse. Osteoarthritis Cartilage 18, S17–S23 (2010).

30. Krenn, V. et al. Synovitis score: discrimination between chronic low-grade and high-grade synovitis. Histopathology 49, 358–364 (2006).

31. Shapouri-Moghaddam, A. et al. Macrophage plasticity, polarization, and function in health and disease. J. Cell. Physiol. 233, 6425–6440 (2018).

32. Funes, S. C., Rios, M., Escobar-Vera, J. & Kalergis, A. M. Implications of macrophage polarization in autoimmunity. Immunology 154, 186–195 (2018).

33. Lumeng, C. N., Bodzin, J. L. & Saltiel, A. R. Obesity induces a phenotypic switch in adipose tissue macrophage polarization. J. Clin. Invest. 117, 175–184 (2007).

34. Kijowski, R., Demehri, S., Roemer, F. & Guermazi, A. Osteoarthritis year in review 2019: imaging. Osteoarthritis Cartilage 28, 285–295 (2020).

35. Chatterjee, P. et al. Adipocyte fetuin-A contributes to macrophage migration into adipose tissue and polarization of macrophages. J. Biol. Chem. 288, 28324–28330 (2013).

36. Oishi, Y. & Manabe, I. Macrophages in inflammation, repair and regeneration. Int. Immunol. 30, 511–528 (2018).

37. Feehan, K. T. & Gilroy, D. W. Is Resolution the End of Inflammation? Trends Mol.Med. 25, 198–214 (2019).

38. Proto, J. D. et al. Regulatory T Cells Promote Macrophage Efferocytosis during Inflammation Resolution. Immunity 49, 666–677.e6 (2018).

39. Yurdagul, A. J. et al. Macrophage Metabolism of Apoptotic Cell-Derived Arginine Promotes Continual Efferocytosis and Resolution of Injury. Cell Metab. 31, 518–533.e10 (2020).

40. Kourtzelis, I. et al. DEL-1 promotes macrophage efferocytosis and clearance of inflammation. Nat. Immunol. 20, 40–49 (2019).

41. Zhen, Y., Lee, I. J., Finkelman, F. D. & Shao, W.-H. Targeted inhibition of Axl receptor tyrosine kinase ameliorates anti-GBM-induced lupus-like nephritis. J. Autoimmun. 93, 37–44 (2018).

42. Szondy, Z., Garabuczi, E., Joós, G., Tsay, G. J. & Sarang, Z. Impaired clearance of apoptotic cells in chronic inflammatory diseases: therapeutic implications. Front. Immunol. 5, 354 (2014).

